# Developmental trajectory of pre-hematopoietic stem cell formation from endothelium

**DOI:** 10.1101/848846

**Authors:** Qin Zhu, Peng Gao, Joanna Tober, Laura Bennett, Changya Chen, Yasin Uzun, Yan Li, Melanie Mumau, Wenbao Yu, Bing He, Nancy A. Speck, Kai Tan

**Author notes:** These authors contributed equally to this work (Qin Zhu, Peng Gao, Joanna Tober).

## Abstract

Hematopoietic stem and progenitor cells (HSPCs) differentiate from hemogenic endothelial (HE) cells through an endothelial to hematopoietic cell transition (EHT). Newly formed HSPCs accumulate in intra-arterial clusters (IACs) before colonizing the fetal liver. To examine the cell and molecular transitions during the EHT, and the heterogeneity of HSPCs within IACs, we profiled ∼37,000 cells from the caudal arteries of embryonic day 9.5 (E9.5) to E11.5 mouse embryos by single-cell transcriptome and chromatin accessibility sequencing. We identified an intermediate developmental stage prior to HE that we termed pre-HE, characterized by increased accessibility of chromatin enriched for SOX, FOX, GATA, and SMAD motifs. A developmental bottleneck separates pre-HE from HE, with RUNX1 dosage regulating the efficiency of the pre-HE to HE transition. Distinct developmental trajectories within IAC cells result in two populations of CD45^+^ HSPCs; an initial wave of lympho-myeloid-biased progenitors, followed by precursors of hematopoietic stem cells (pre-HSCs).

## Introduction

Hematopoietic ontogeny involves multiple “waves” in which HSPCs with different potentials differentiate from HE cells. HE cells in the yolk sac (YS) differentiate into committed erythro-myeloid (EMP) and lymphoid progenitors, and the caudal arteries produce lymphoid progenitors and pre-HSCs (Auerbach et al., 1996; Hadland and Yoshimoto, 2018). YS hematopoiesis can be recapitulated in embryonic stem (ES) cell cultures, where the molecular events are well-described (Goode et al., 2016; Vanhee et al., 2015). Although groundbreaking studies described the transcriptomes of HE and pre-HSCs in the major caudal artery, the dorsal aorta, at single cell resolution (Baron et al., 2018; Zeng et al., 2019; Zhou et al., 2016), these analyses did not examine the distribution of cells along the trajectory from E to pre-HSCs, or the chromatin landscapes of cells along the trajectory, due to the limited number of cells sequenced. To gain insights into the molecular mechanisms mediating the differentiation of arterial E cells into IACs and to examine heterogeneity within IACs, we utilized the 10x Genomics platform and performed single-cell RNA sequencing (scRNA-Seq) and single-cell assay for transposase-accessible chromatin sequencing (scATAC-Seq). Our data reveal a continuous trajectory from E to IAC cells, previously undefined transitional cell populations along the trajectory, the pathways and transcription factors active in these cells, and describe the molecular heterogeneity of IAC cells.

## Results

### scRNA-sequencing reveals a continuous trajectory from endothelial cells to IAC cells

Our strategy was to analyze all cells along the trajectory in a single sample to determine their distribution between different transcriptional states, and combine that with analyses of purified sub-populations to make accurate cell assignments and obtain additional coverage of rare cells. We captured the entire trajectory by purifying a population containing all E, HE, and IAC cells (E+HE+IAC) from E9.5 and E10.5 embryos using a combination of endothelial markers (Fig. 1a,b; Supplementary Fig. 1). We then purified subpopulations of HE and E cells based on expression of green fluorescent protein (GFP) from the *Runx1* locus (Lorsbach et al., 2004) (Fig. 1c). We confirmed that only HE cells were capable of producing hematopoietic cells *ex vivo* (Fig. 1d). Kit^hi^ IAC cells were excluded in the sorts, therefore HE and E cells were negligibly contaminated with HSPCs (Fig. 1e). We purified IAC cells from E10.5 and E11.5 embryos using antibodies recognizing endothelial markers and Kit (Fig. 1f), E9.5 yolk sac EMPs, and E14.5 fetal liver HSCs (FL-HSCs) (Supplementary Fig. 1).

**Figure 1.**
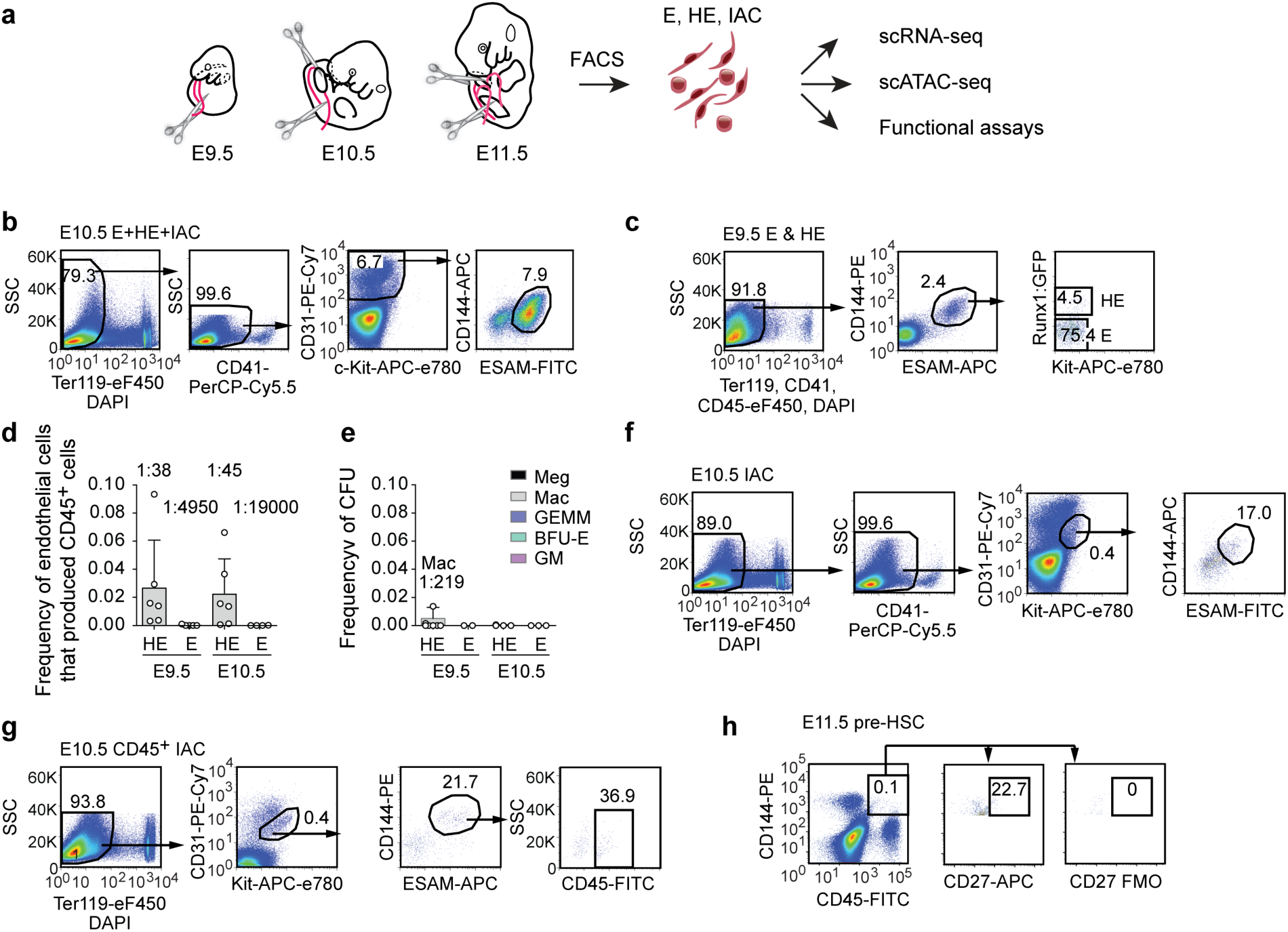
Experimental design, sorting and functional characterization of cell populations. a. The caudal part of embryos were isolated (boundaries are illustrated with scissors), then organs and gut tube removed. Vitelline and umbilical arteries (VU) were isolated and included in the sample. The tissue was dissociated and cells were isolated by FACS, then analyzed by scRNA-seq, scATAT-seq, or in functional assays. All cell populations purified and sequenced are listed in Supplemental Table 1, and additional sort plots are in Supplementary Fig. 1. b. Representative scatter plots for purification of E+HE+IAC cells (Ter119^-^CD41^med/-^ CD31^+^CD144^+^ESAM^+^) from the caudal half of E10.5 embryos. Kit was not used in the sorts in order to capture cells throughout the trajectory, including IAC cells. Numbers on the x and y-axes are indicated on the first FACS plot on the left, and unless changed are not depicted on plots to the right of the preceding plot. See Supplementary Fig. 1a for E9.5 E+HE+IAC purification. c. Representative scatter plots for purification of E (Ter119^-^CD41^-^CD45^-^ CD31^+^CD144^+^ESAM^+^Kit^-^GFP^-^) and HE (Ter119^-^CD41^-^CD45^-^CD31^+^CD144^+^ESAM^+^Kit^lo/-^ GFP^+^) cells from E9.5 RUNX1:GFP embryos(Lorsbach et al., 2004). See Supplementary Fig. 1b for E10.5 E and HE purifications. d. Frequency of purified endothelial cells from E9.5 and E10.5 embryos that gave rise to CD45^+^cells when cultured in a limiting dilution assay on OP9 stromal cells. Frequencies are indicated on top of the bars. HE, Runx1:GFP^+^ endothelial cells, purified as described in panel c; E, Runx1:GFP^-^ endothelial cells. Error bars, mean ± standard deviation (SD); n = 5-6 experiments. Frequencies were calculated using ELDA software(Hu and Smyth, 2009). e. Colony forming units (CFU) representing frequency of contaminating committed HSPCs in purified HE and E populations. Meg, megakaryocyte; Mac, macrophage; GEMM, granulocyte/erythroid/monocyte/megakaryocyte; BFU-E, burst forming unit-erythroid; GM, granulocyte/monocyte. Error bars, mean ± SD; n = 2-3 experiments. f. Representative scatter plots of E10.5 IAC cells (Ter119^-^CD41^med/-^ CD31^+^CD144^+^ESAM^+^Kit^+^). See Supplementary Fig. 1c for E11.5 IAC cell purification. g. Representative scatter plots for purification of E10.5 CD45^+^ IAC cells containing lympho-myeloid biased progenitors (Ter119^-^CD41^med/-^CD31^+^CD144^+^ESAM^+^Kit^+^CD45^+^). h. Representative scatter plots for purification of E11.5 CD45^+^CD27^+^CD144^+^ IAC cells containing type II pre-HSCs (E11.5 pre-HSC).

Summary statistics for collected cell populations are in Fig. 2a and Supplementary Table 1. We used uniform manifold approximation and projection (UMAP) to reduce the data dimension (Becht et al., 2018). After filtering out non-endothelial and non-hematopoietic cells, UMAP of the combined datasets shows a continuous trajectory from E to IAC cells (Fig. 2b,c, Supplementary Fig. 2). E14.5 FL-HSCs are disconnected from this trajectory, therefore, are more distantly related (Fig. 2b). Two streams of E cells expressing the arterial marker *Efnb2* converge to form a stem leading to HE and IACs (Fig. 2d). Analyses of E10.5 E+HE+IAC cells manually separated into VU arteries and DA demonstrated that VU cells contribute to one of these streams and DA to both streams (Fig. 2e). The E+HE+IAC samples, which demonstrate the distribution of cells at various stages, show that at E9.5, IAC cells constitute only 0.5% of the population, but at E10.5 the fraction of IAC cells expands by 7 fold, representing 3.5% of the population, consistent with histological analyses of the temporal appearance of IACs in embryos at those two stages (Garcia-Porrero et al., 1995; Yokomizo and Dzierzak, 2010) (Fig. 2c,f).

**Figure 2.**
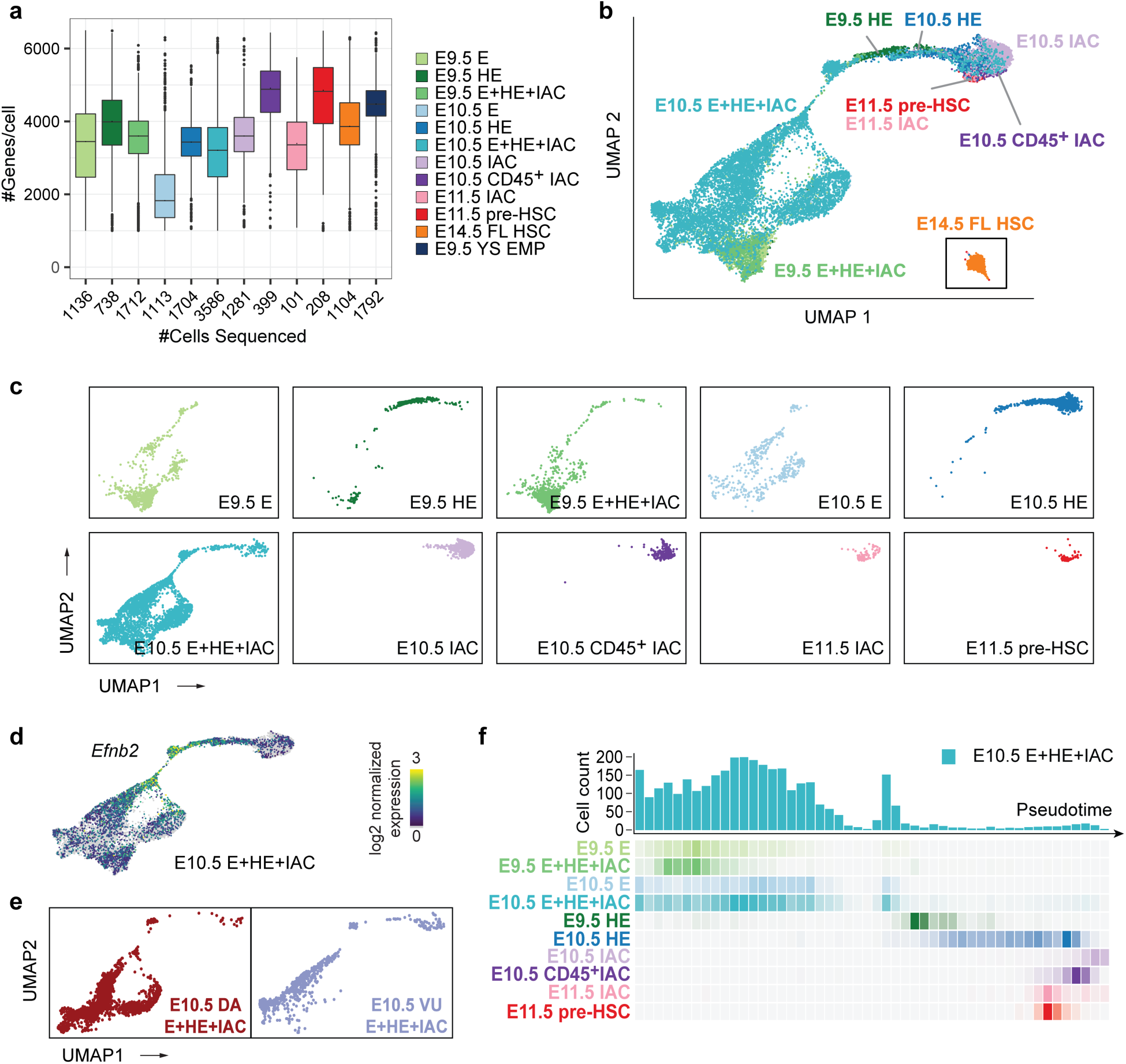
Overview of single cell RNA-Seq data. a. The number of cells sequenced (x-axis) and genes per cell detected for representative samples. b. UMAP of continuous EHT trajectory and FL-HSCs, with selected cell populations labeled. c. Distribution of cells from each dataset in the UMAP reflecting EHT trajectory. d. UMAP illustrating the two streams of arterial E cells expressing high levels of *Efnb2* that converge to form the stem leading to HE and IACs. e. E+HE+IAC cells separately purified from the vitelline and umbilical (VU) arteries, and from the dorsal aorta (DA) within the caudal half of the embryo, highlighted on the global UMAP plot. f. Cell count along the pseudotime trajectory. Bar graph quantifies results from a single sort of E10.5 E+HE+IAC cells; heat maps below the graphs show distribution of cells in all sorted cell populations.

Louvain clustering identified 7 distinct populations in the combined dataset, and separated the two streams of arterial E cells into distinct clusters (Fig. 3a). One cluster of cells, Wnt^hi^ E containing only DA cells, expresses high levels of Wnt target genes. The second cluster, Wnt^lo^ E containing both DA and UV cells, expresses lower levels of Wnt target genes. Wnt^hi^ E and Wnt^lo^ E converge to form a distinct cluster we term conflux E. Pseudo-time-ordered Wnt^hi^ and Wnt^hi^ E cells exhibit gradual decreases in transcriptome similarity, followed by a sharp increase as cells converge in conflux E (Fig. 3b,c). The increase in transcriptome similarity is driven by the loss of cluster-specific gene expression and increased transcript levels from late-stage-specific genes (Fig. 3d,e). For example, Wnt^hi^ E expresses high levels of Wnt target genes including *Foxq1,* while Wnt^lo^ E is characterized by high levels of *Hpgd* transcripts (Fig. 3e). Both Wnt target genes and *Hpgd* are subsequently significantly down-regulated in conflux E. Conflux E is characterized by elevated Notch signaling, indicated, for example, by increased expression of the Notch ligands *Jag1* and *Dll4* and the transcription factor *Hey2* (Fig. 3e,f; Supplementary Figure 2c). Additional pathways activated during the convergence of Wnt^hi^ and Wnt^hi^ E into conflux E include pathways regulating cell shape and motility (“assembly of collagen fibrils”, “laminin interactions”, “gap junction trafficking”), lipid metabolism (“triglyceride biosynthesis”) and processes important in hematopoietic cells (“interleukin-6 family signaling”, “MAPK signaling for integrins”) (Fig. 3f).

**Figure 3.**
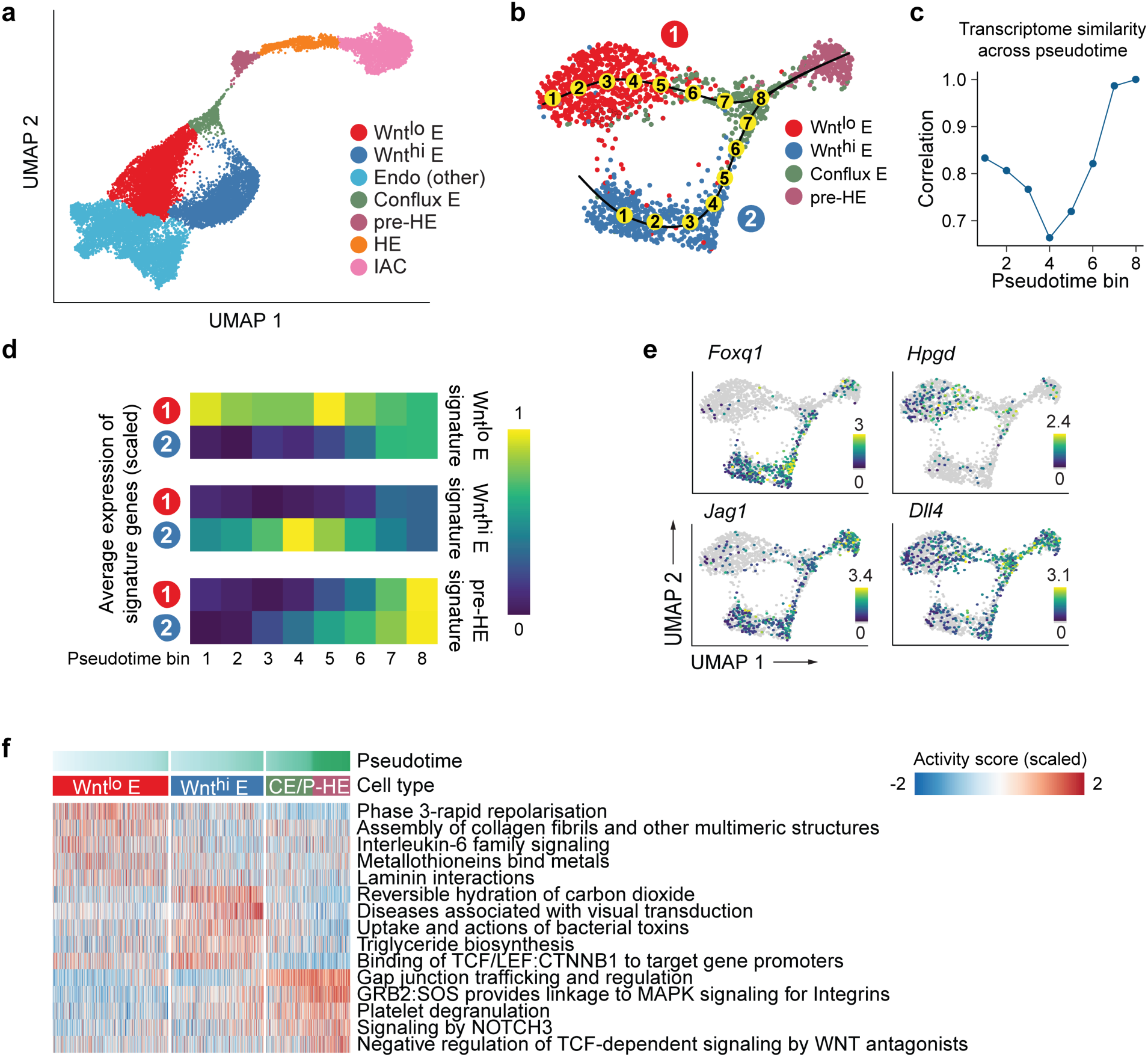
Two streams of endothelial cells converge before hemogenic endothelium. a. UMAP of EHT trajectory (from Fig. 2b, with FL-HSC removed) showing 7 clusters identified by Louvain clustering and differential expression analysis. b. Zoom-in UMAP highlighting the two streams of endothelial cells (1: Wnt^lo^ E; 2: Wnt^hi^ E) converging to conflux E. Numbers in yellow circles represent pseudotime bins up to the point of convergence. c. Pearson correlation between average gene expression in the two endothelial streams over pseudotime bins. d. Average expression of Wnt^lo^ E, Wnt^hi^ E, and pre-HE-specific genes over pseudotime. Differentially expressed genes were derived by pairwise expression analysis between Wnt^lo^ E and Wnt^hi^ E. Pre-HE specific genes were derived by comparing pre-HE with Wnt^lo^ plus Wnt^hi^ E. e. Expression of Wnt target *Foxq1* and *Hpgd* decrease in conflux E, although *Foxq1* expression is regained in a subset of pre-HE cells. Notch ligand *Jag1* is up-regulated in Wnt^hi^ E, while *Dll4* is expressed in both Wnt^lo^ and Wnt^hi^ E. Expression of *Jag1* and *Dll4* is elevated post convergence. f. Heatmap showing stream-specific Reactome pathway activity over pseudotime. Cells were grouped into Wnt^lo^ E, Wnt^hi^ E and CE/PHE (Conflux E or pre-HE) and ordered using pseudotime. AUCell package (Aibar et al., 2017) was used to compute a pathway activity score for each cell. One vs the rest Student *t* test were used to derive group-specific pathways and the top 5 most significant pathways plotted.

### Runx1 regulates progression through a developmental bottleneck between pre-HE and HE

HE cells are characterized by high levels of *Runx1* and *Gfi1*, and IAC cells at the far end of the trajectory are defined by expression of the pan-hematopoietic marker gene *Ptprc* (encoding CD45) (Fig. 4a,b; Supplementary Fig. 2c). Between conflux E and HE is a distinct cluster of endothelial cells that we named pre-HE. UMAP and pseudotime trajectories of E10.5 E+HE+IAC reveal an accumulation of pre-HE cells, suggesting a bottleneck between pre-HE and HE (Fig. 2f, 4a). *Gfi1*, a direct RUNX1 target that participates in extinguishing endothelial fate (Lancrin et al., 2012), shows elevated expression immediately after cells pass through the bottleneck and become HE, while high levels of *Sox17,* the Notch target *Hey2,* and the arterial marker *Cd44* are found in pre-bottleneck populations including conflux E and pre-HE (Fig. 4b). To provide further evidence for the bottleneck, we utilized Velocyto, which infers directionality of differentiation by modeling dynamics of unspliced versus spliced RNAs when a gene is up or down-regulated (La Manno et al., 2018). Velocyto showed a marked decrease in RNA velocity in pre-HE cells, and many appear to revert to an earlier transcriptome state, suggesting a differentiation barrier restricting their progression towards HE (Fig. 4c). Once pre-HE cells transit to HE, however, they smoothly differentiate to IAC cells. Several pathways known to promote *Runx1* expression and HSPC formation are upregulated in pre-HE, including Notch, tumor necrosis factor, fluid shear stress, cytokine signaling, and synthesis of eicosanoids, vitamins, and sterols (Adamo et al., 2009; Cortes et al., 2016; Espin-Palazon et al., 2014; Gu et al., 2019; Kim et al., 2014; Li et al., 2015; North et al., 2009; North et al., 2007), suggesting these pathways are important in pre-HE (Fig. 4d, Supplementary Fig. 3a, Supplementary Table 2). Once cells transition to HE, RUNX1 plays a predominant role.

**Figure 4.**
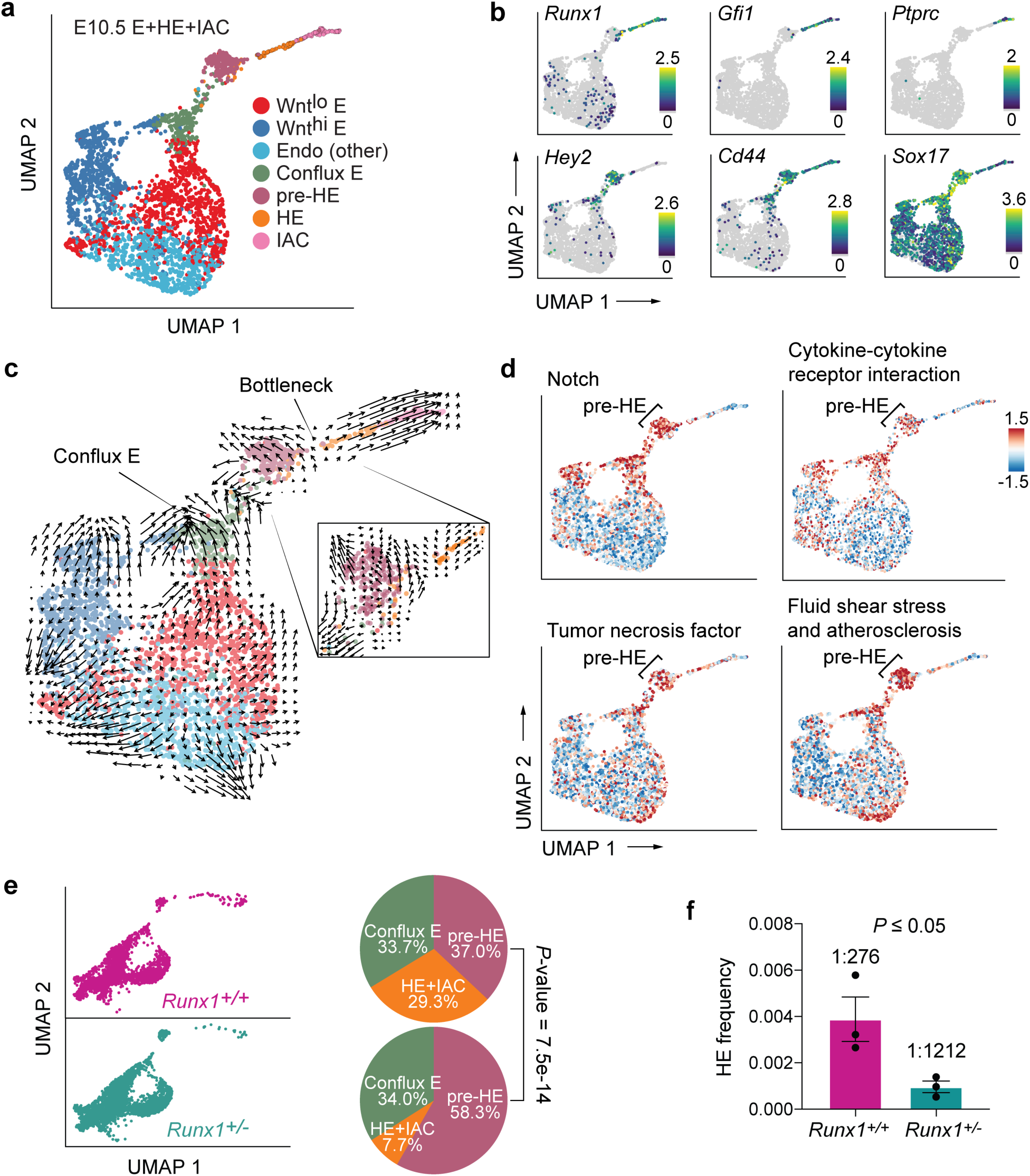
Developmental bottleneck between pre-HE and HE cells. a. UMAP of E10.5 E+HE+IAC cells showing 7 clusters from Fig. 3a. b. Expression of key markers of clusters, including *Hey2* in conflux E and pre-HE, *Cd44* in conflux E, pre-HE, HE and IACs, *Ptprc* in IACs, *Gfi1* and *Runx1* in HE and IACs, and high levels of *Sox17* in conflux E and pre-HE, with downregulation in HE. Note *Runx1* is expressed at low levels in a subset of Wnt^lo^ E, Wnt^hi^ E, and Endo (other) cells. c. Velocyto analysis revealing different differentiation dynamics along the EHT in E10.5 E+HE+IAC cells. To the right is a zoom-in velocity of pre-HE cells that have accumulated at the bottleneck between pre-HE and HE. d. Activity of pathways from Kyoto Encyclopedia of Genes and Genomes (KEGG) database, computed for each cell using the AUCell method (Aibar et al., 2017). e. UMAP of E+HE+IAC cells from E10.5 *Runx1^+/+^* and *Runx1^+/-^* littermates. Pie charts to the right depict the distribution of cells between conflux E, pre-HE, and combined HE+IAC populations in E10.5 *Runx1^+/+^* and *Runx1^+/-^* littermates. HE and IAC cells were combined to increase the statistical power. *P*-values indicate significant differences in the distributions of cells in pre-HE and HE+IAC in *Runx1^+/+^* versus *Runx1^+/-^* samples based on proportion test. f. Limiting dilution assay to determine the frequency of HE in the CD44^+^ fraction of E+HE+IAC cells isolated from E10.5 embryos (see Supplementary Fig. 1g for FACS plots). Shown are frequencies of cells that yielded hematopoietic cells (B220^+^, CD19^+^, Mac1^+^, Gr1^+^, and/or CD41 and CD45) *ex vivo.* Frequencies were calculated by ELDA(Hu and Smyth, 2009). Data represent three independent cell purifications and limiting dilution assays (mean ± SD, unpaired two-tailed Student’s *t-*test).

*Runx1* expression is upregulated in approximately 7% of pre-HE cells suggesting that RUNX1 levels regulate passage through the bottleneck (Fig. 4b). To test this hypothesis, we compared the distribution of cells between conflux E, pre-HE, and HE+IAC in *Runx1^+/-^* and *Runx1*^+/+^ littermates. We observed a 74% reduction in the proportion of HE and IAC cells in E10.5 *Runx1^+/-^* compared to *Runx1*^+/+^ embryos, and a commensurate 58% increase in pre-HE, consistent with the hypothesis that Runx1 levels regulate transit through the bottleneck (Fig. 4e). To confirm that RUNX1 haploinsufficiency reduces functional HE cells, we utilized the arterial marker CD44 (Wheatley et al., 1993) in conjunction with other endothelial markers to purify cells enriched for conflux E, pre-HE, HE, and IAC (Supplementary Fig. 1g) and performed limiting dilution assays. RUNX1 haploinsufficiency reduced the frequency of functional HE cells by 77% (Fig. 4f), equivalent to the reduction in UMAP plots (Fig. 4e), confirming that RUNX1 levels regulate the number of pre-HE cells that transit through the bottleneck to become HE cells.

### scATAC-Seq identifies putative *Runx1* enhancers and transcription factor motifs that gain accessibility in pre-HE

To identify signals that may activate *Runx1* expression in pre-HE, we performed paired scRNA-Seq and scATAC-Seq on E10.5 CD44^+^ E+HE+IAC cells to identify *Runx1* enhancers and the stages they are accessible. High quality open chromatin profiles were obtained for 1670 cells, covering various cell types from E to IAC (Supplementary Fig. 4). We were unable to separate HE from pre-HE cells due to the small representation of HE cells in the sample, therefore HE cells segregated with pre-HE. We devised a computational approach that aligns scATAC-Seq and scRNA-Seq data (Fig. 5a, Supplementary Fig. 5,6) and subsequently linked enhancers with their target promoters (Fig. 5b). Accuracy of our method was benchmarked using known hematopoietic and endothelial enhancers (Fig. 5c-d). We applied chromVar (Schep et al., 2017) to assess differential transcription factor (TF) binding patterns along the EHT trajectory (Fig. 5e). Results show strong correlation with the TF expression patterns and are consistent with pathway analyses from scRNA-seq data. For example, strong TCF/LEF binding activity was detected in Wnt^hi^ arterial E that abruptly decreased in conflux E (Fig. 5e,f). SOX and FOX binding sites are mostly open in conflux E and pre-HE. Binding sites for a large group of TFs had increased accessibility beginning at the pre-HE stage, including HES1, GATA, SMAD, and TFs that regulate HSC homeostasis (MECOM, EGR1, YY1) (Goyama et al., 2008; Lu et al., 2018; Min et al., 2008), the latter group suggesting that an HSC-specific transcriptional program may initiate at the pre-HE stage.

**Figure 5.**
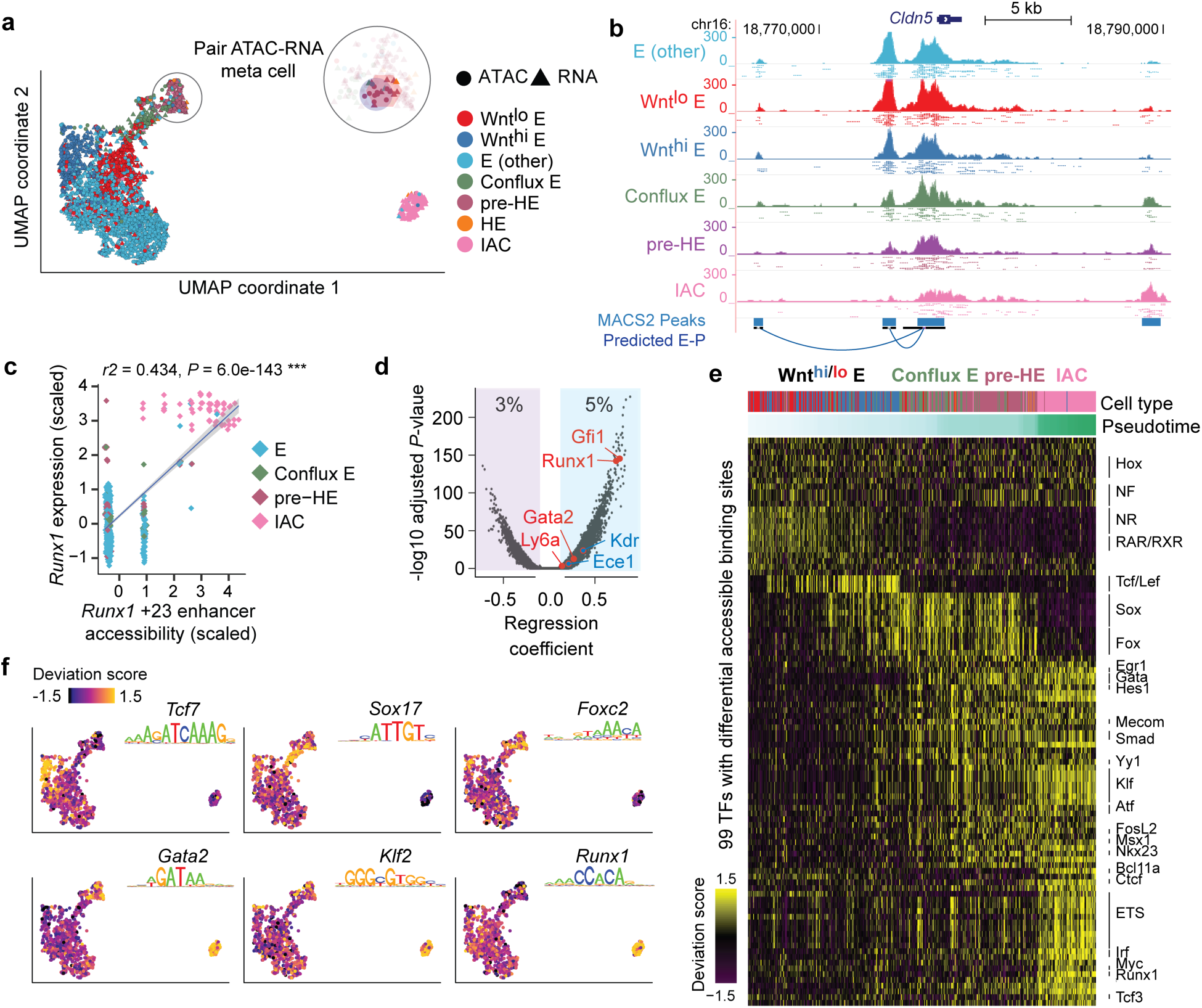
Joint scRNA-Seq and ATAC-Seq analysis of bottleneck populations. a. UMAP of 1637 cells from scRNA-Seq and 1186 cells from scATAC-Seq, aligned using Seurat algorithm with a custom defined gene-peak correspondence matrix (see Methods). The number of HE cells was too few to be resolved by UMAP, and clustered with pre-HE. To gain enough statistical power for predicting E-P, we pooled reads from 10 nearest neighbors as “meta cells”, and paired scATAC meta cell to nearby scRNA meta cell. Additional details can be found in the Methods section. b. UCSC genome browser tracks showing open chromatin signal of *Cldn5* promoter and its predicted enhancers. Dots below each aggregated signal track presents signal from 50 sampled cells of each type. c. Linear regression shows high correlation between *Runx1* +23 enhancer chromatin accessibility and *Runx1* expression levels (z-score transformed). Each point represents a paired ATAC-RNA meta cell in panel a, with pooled RNA expression on the y-axis and pooled enhancer accessibility on the x-axis. d. Linear regression result for each expressed gene against scATAC-Seq peaks 200k bp upstream and downstream of its transcription start site (TSS). We recapitulated the majority of known enhancer-promoter interactions (E-Ps) that function during EHT, with the *Runx1* +23 enhancer (Nottingham et al., 2007) and *Gfi1* enhancer (Wilson et al., 2010) among the top predictions. e. TF binding patterns among called scATAC-Seq peaks assessed using chromVar (Schep et al., 2017), which defines a deviation score reflecting the accessibility change at binding sites of each TF across all cells. Binding sites were determined using DNA motif scan on the called enhancers, which does not discriminate TFs in the same family with very similar motifs. Top significant TFs based on Mann-Whitney U test are plotted for each stage. f. ChromVar deviation score for selected TF motifs plotted on the UMAP, showing specific binding pattern for Tcf7 in Wnt^hi^ E, Sox17 in conflux E, Foxc2 in pre-HE, Gata2 and Klf2 in both pre-HE and IAC. Runx1 binding sites are highly accessible post bottleneck, but also exhibit medium to high level of chromatin accessibility in some early stage cells.

*Runx1* contains two promoters, an upstream P1 promoter that is first utilized in committed HSPCs, and a more proximal P2 promoter that is active in HE and HSPCs (Sroczynska et al., 2009). Consistent with this, P1 first becomes accessible in IAC cells, whereas P2 is accessible in all endothelial cells including pre-HE (Fig. 6a), which may permit or contribute to the stochastic *Runx1* expression observed in a subset of endothelial cells (Fig. 4b). Using our computational approach, we predicted 27 enhancer-promoter (E-P) interactions, which recapitulate 11 out of 22 previously identified E-Ps based on chromosome conformation capture assays (Chen et al., 2019; Marsman et al., 2017) (Fig. 6a, also see Methods). All of the predicted enhancers exhibit higher co-accessibility with P1 compared to P2, therefore only E-Ps to P1 are indicated. Significance plot of the predicted E-Ps reveals several enhancers whose chromatin openness is significantly correlated with *Runx1* expression, including the *Runx1* +23 enhancer (Fig. 6b). Several of the predicted enhancers exhibit stage-specific co-accessibility with the P1 promoter (Fig. 6c). Interestingly, one previously functionally-validated enhancer located 371 kb upstream of *Runx1* P1 (Marsman et al., 2017) was accessible only in pre-HE and IACs, and not in other endothelial cell populations, suggesting it could initiate the opening of P1 in pre-HE, and contribute to the activity of P1 in IAC cells (Fig. 6a,c). The ATAC-seq signal encompassing the −371 enhancer begins to increase in conflux E cells and reaches a maximum in pre-HE cells (Fig. 6a,d). This change in accessibility in pre-HE coincides with the activation of *Runx1* expression in a subset of pre-HE cells (Fig. 6d). However, unlike the +23 enhancer, the chromatin accessibility of the −371 enhancer subsequently decreases in IAC cells and is no longer open in FL-HSCs (Fig. 6a). The −371 enhancer contains GATA, STAT, and JUN motifs, indicating that GATA2 and cytokine and/or inflammatory signaling may contribute to the opening of this enhancer in pre-HE (Fig. 6a). An independent co-expression analysis based on the scRNA-Seq data reveals that these factors form a co-expression gene module that precedes and correlates with *Runx1* expression (Figure 6e), suggesting they may cooperatively regulate *Runx1* expression. Notably, neither the −371 nor +23 enhancers contain SOX motifs, which are recognized by a repressor of Runx1 expression, Sox17 (Bos et al., 2015). Other TF motifs enriched in the 27 called *Runx1* enhancers, all of which are predicted to interact with P1, include ETS, FOX, SOX, KLF/SP, RUNX, and SMAD, which are recognized by TFs with well-documented roles in HSPC formation (Blank and Karlsson, 2011; Gilmour et al., 2019; Menegatti et al., 2019).

**Figure 6.**
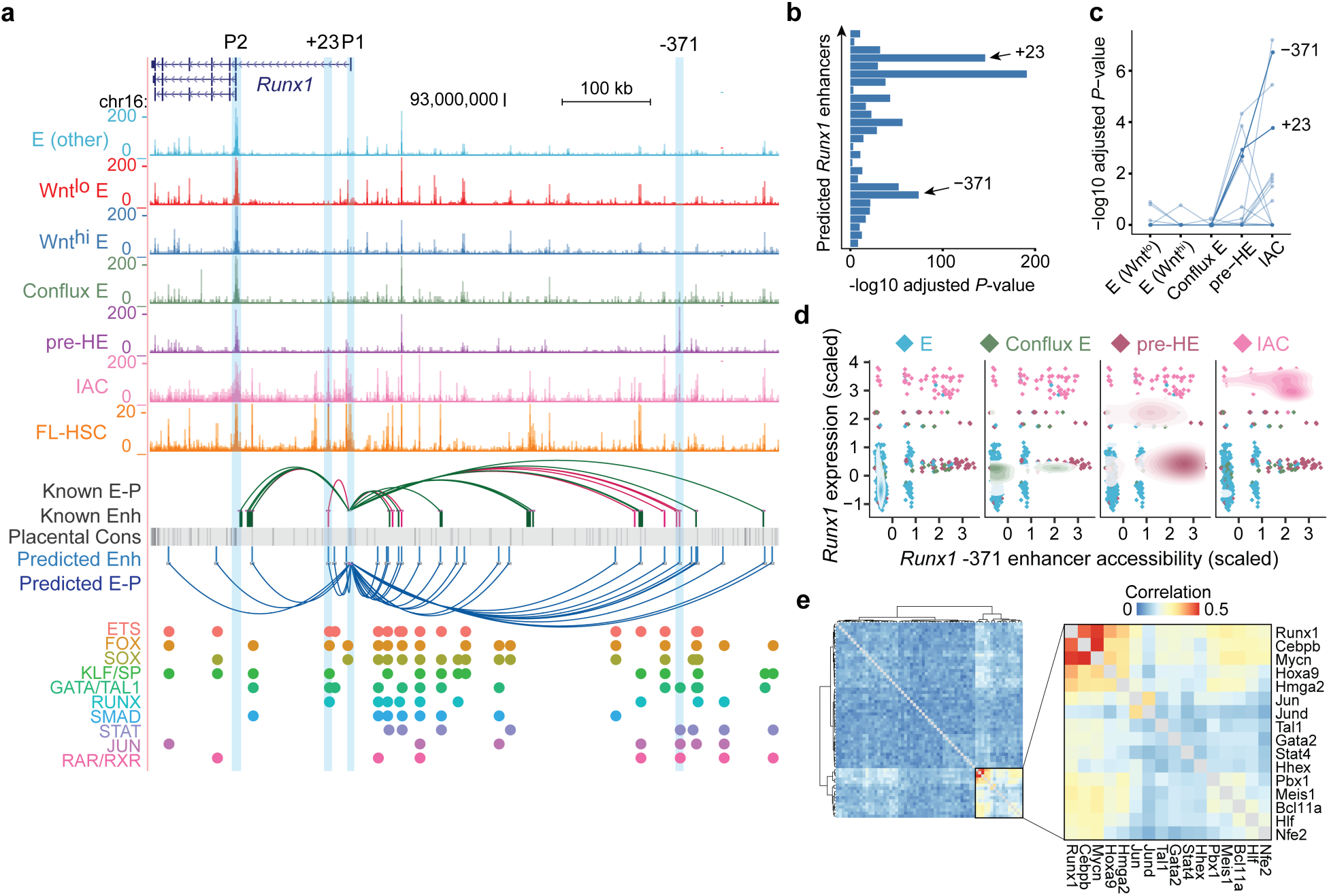
Developmental-stage-specific enhancers of *Runx1*. a. UCSC genome browser tracks showing open chromatin signal for each of the populations. Tracks from E to IAC are cumulative scATAC-Seq signals (per-base unique fragment coverage) normalized by the number of cells in that population. Tracks for FL-HSC are bulk ATAC-Seq data from Chen, C. *et al* (Chen et al., 2019). Experimentally validated enhancers and E-Ps from Marsman *et al*. (Marsman et al., 2017) are shown in magenta. Enhancers and E-P links from Chen, C. *et al.* (Chen et al., 2019) are shown in dark green. E-P links were inferred based on linear regression on paired scRNA-scATAC meta cells (see Methods). Placental mammal conservation by PhastCons score is shown as grey track. For each of the inferred enhancers, we scanned for known motifs from CIS-BP database and grouped TFs from the same family having similar motifs. Motif hits of several previously reported early hematopoietic TFs are highlighted below the track. b. Distribution of linear regression P-values for predicted *Runx1* enhancers. Highly significant peaks include the validated +23 and −371 enhancers. The most significant peak is ∼3.6 kb downstream of P1. c. Co-accessibility of *Runx1* P1 promoter and its predicted linked enhancers in each cell type. P-values for co-accessibility in each cell type were computed using Fisher’s exact test with multiple testing correction. d. Stage-specific chromatin accessibility of *Runx1* −371 enhancer and *Runx1* expression levels (z-score transformed). Each point in the scatter plot represents a paired ATAC-RNA meta cell in Fig. 5a, with pooled RNA expression on the y-axis and pooled enhancer accessibility on the x-axis. A 2-dimensional density plot is superimposed on the scatter plot. e. Co-expression of transcription factors that have binding motifs at *Runx1* enhancers and whose expression precedes *Runx1*. Correlations were computed using gene expression matrix including conflux E, pre-HE and HE cells. TFs with Pearson correlation with Runx1 < 0.05 were removed. Hierarchical clustering was performed on the correlation matrix and a strong TF co-expression module was highlighted.

### Two waves of HSPCs form in the IACs

We also examined the transition of HE to IAC cells and the composition of IAC cells. Principal component analysis (PCA) depicts a sharp U-turn as HE differentiates into IAC cells, reflective of a marked decrease in arterial endothelial gene expression and activation of hematopoietic genes (Fig. 7a). For example, the arterial E-specific gene *Gja5* is primarily expressed on the HE side of the trajectory, while expression of *Spn* (encoding CD43), *Ptprc* (encoding CD45) and the Rho GTPase *Rac2* rapidly increases in IAC cells (Fig. 7b, Supplementary Fig. 3b,c). Transient expression of the chromatin remodeling protein *Nupr1* occurs at the U-turn, while *Hey1* and *Sox17* transcripts significantly diminish as IAC cells mature (Fig. 7b, Supplementary Fig. 3c).

**Figure 7.**
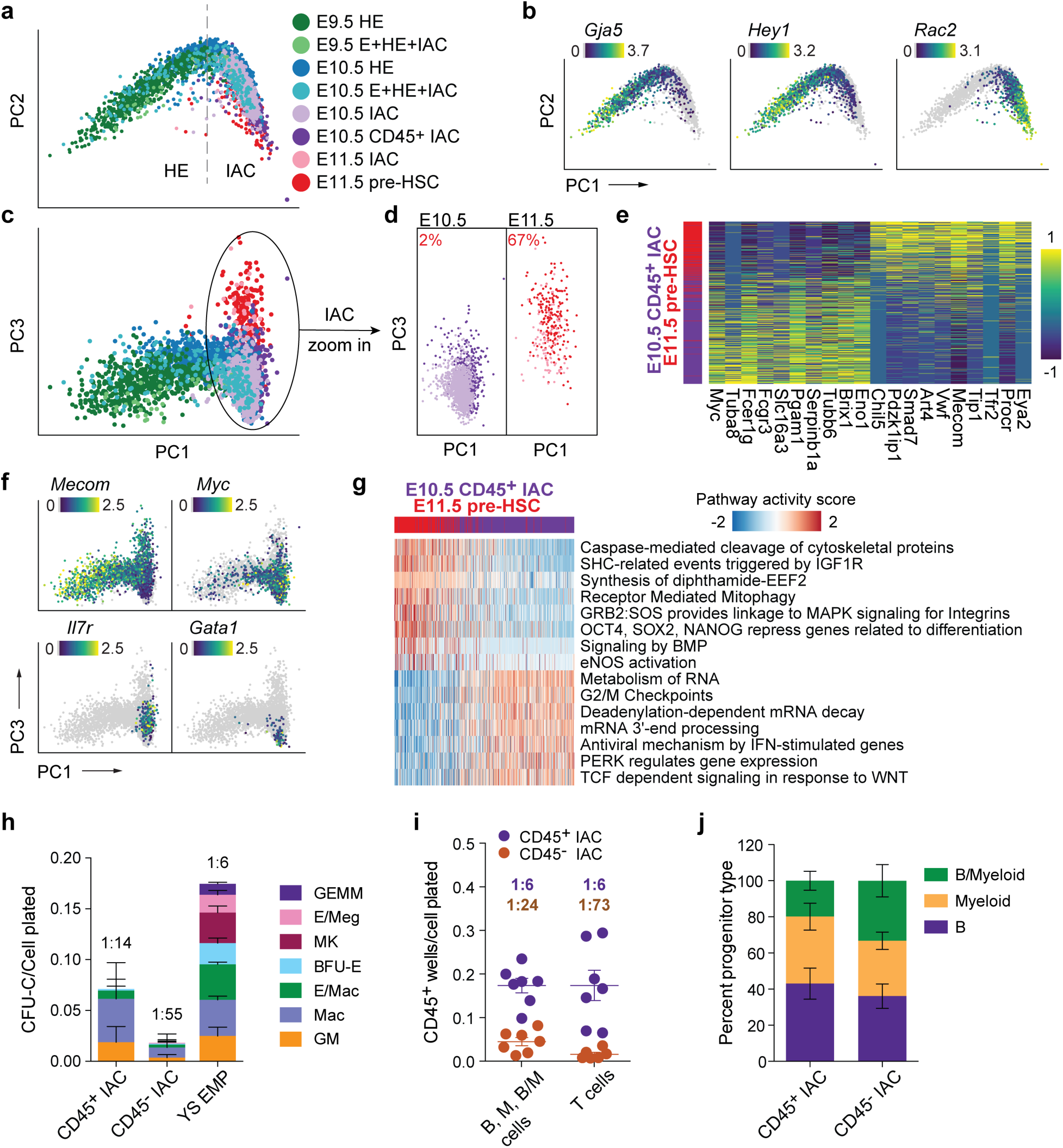
Two waves of CD45^+^ HSPCs in IAC cells. a. PCA plot of a subset of data containing IACs, illustrating the trajectory of IAC differentiation from HE along the PC1 axis. b. Expression of *Gja5*, *Hey1*, and *Rac2* illustrating the maturation of IAC cells along the trajectory. c. PCA plot showing the separation of E10.5 and E11.5 IAC cells along the PC3 axis. d. E10.5 and E11.5 IAC cells, E10.5 CD45^+^ IAC cells, and E11.5 pre-HSCs plotted separately to visualize their relative distribution along the PC3 axis. A k-nearest-neighbor classifier (k=3 with PC1-10 as feature input) was trained using E10.5 CD45^+^ IAC cells and E11.5 pre-HSCs to determine the fraction of pre-HSCs (labeled in red) in E10.5 and E11.5 IAC cells. e. Heat map showing top differentially expressed genes in E10.5 CD45^+^ IAC cells versus E11.5 pre-HSCs. f. Preferential expression of *Mecom* in E11.5 pre-HSCs and IAC cells, versus *Myc, Il7r*, and *Gata1* in E10.5 CD45^+^ IAC enriched for lympho-myeloid-biased progenitors and in E10.5 IAC cells. g. Reactome pathway analysis comparing E11.5 pre-HSC and E10.5 CD45^+^ IAC cells. Color indicates pathway activity score computed using the AUCell package (Aibar et al., 2017). h. Methylcellulose (colony forming unit-culture, CFU-C) assay performed in the presence of stem cell factor (SCF), interleukin 3 (IL3), IL6, and erythropoietin (EPO) to measure the frequency of committed erythroid and myeloid progenitors in E10.5 CD45^+^ IAC cells, CD45^-^ IAC cells, and E9.5 yolk sac EMPs. BFU-E, burst forming unit-erythroid; GM, granulocyte/macrophage; Mac, macrophage; MK, megakaryocyte; GEMM, granulocyte/erythroid/monocyte/megakaryocyte. Error bars; mean ± SD. Frequencies of total progenitors are indicated above the bars. n = 3 experiments. i. Limiting dilution assays on OP9 stromal cells to determine the frequencies of progenitors in purified E10.5 CD45^+^ IAC and E10.5 CD45^-^ IAC cells yielding B (CD45^+^CD19^+^B220^mid/lo^), myeloid (M) (Gr1^+^Mac1^+^or Gr1^+^Mac1^-^), and B+myeloid (B/M) cells in culture. Also shown are frequencies of progenitors in purified E10.5 CD45^+^ IAC and E10.5 CD45^-^ IAC cells that produced T cells (CD90^+^ CD25^+^) when cultured on OP9 cells expressing the Notch ligand delta like 1. Error bars; mean ± SD. Frequencies of all progenitors are indicated above the bars. n = 7 experiments. j. Percentage of wells at the limiting cell dose containing B, M, or B/M cells from experiments in panel i. n=8 experiments.

IACs contain pre-HSCs that cannot engraft adult mice directly, but can mature *in vivo* or *ex vivo* into adult-repopulating HSCs (Kieusseian et al., 2012; Rybtsov et al., 2016; Rybtsov et al., 2011). Pre-HSCs are classified as type I or II based on CD45 expression; type I are CD45^-^, and the more mature type II are CD45^+^ (Rybtsov et al., 2011). E10.5 IACs contain only type I pre-HSCs, whereas E11.5 IACs contain both type I and II pre-HSCs (Rybtsov et al., 2011). Additionally, multiple progenitors with lymphoid, myeloid, lympho-myeloid, or multi-lineage potential emerge prior to or contemporaneously with pre-HSCs^2^, at least a subset of which are CD45^+^ (Baron et al., 2018; Inlay et al., 2014; Ohmura et al., 1999). We compared E10.5 CD45^+^IAC cells that contain HSC-independent progenitors and lack pre-HSCs to E11.5 CD45^+^CD27^+^CD144^+^ IAC cells enriched for type II pre-HSCs (E11.5 pre-HSCs) (Li et al., 2017; Rybtsov et al., 2011) to determine their developmental relationship (Fig. 1g,h, Supplementary Fig. 1). The two populations bifurcate in the third principal component of PCA plots; specifically, the majority of E10.5 CD45^+^ IAC cells occupy one end of the PC3 axis, and E11.5 pre-HSCs reside on the other end (Fig. 7c,d). E11.5 pre-HSCs demonstrated a high correspondence with previously published data (Zhou et al., 2016) (Supplementary Fig. 7a). We determined the fraction of pre-HSCs in E10.5 and E11.5 IAC cells using a k-nearest-neighbor classifier. About 2% of E10.5 IAC cells were found to be molecularly similar to E11.5 type II pre-HSCs; this fraction of pre-HSCs increases to 67% in E11.5 IAC cells (Figure 7d), consistent with previous limiting dilution assay results demonstrating an increase in functional pre-HSCs between E10.5 and E11.5 (Rybtsov et al., 2016).

The 974 genes more highly expressed in E11.5 pre-HSCs include known markers of pre-HSCs and/or HSCs (*Eya2, Procr, Cd27,* and *Mecom*) (Forsberg et al., 2005; Goyama et al., 2008; Iwasaki et al., 2010; Li et al., 2017; Zhou et al., 2016), while the 877 genes up-regulated in E10.5 CD45^+^ IAC cells include proliferation related genes (*Myc*) and lympho-myeloid associated genes (*Il7r, Fcer1g*) (Fig. 7e,f, Supplementary Table 3). Pathway analysis suggests E11.5 pre-HSCs gain stem-cell specific features such as “OCT4, SOX2, NANOG represses genes related to differentiation”, while pathways associated with E10.5 CD45^+^ IAC cells are associated with cell cycle and/or related to a specific hematopoietic lineage, such as “TCF dependent signaling in response to WNT” (Fig. 7g, Supplementary Table 3). Interestingly, E11.5 pre-HSCs, although sampled 1 day later in development compared to E10.5 CD45^+^ IAC cells, retain many pathways from E/HE stages, such as “Signaling by BMP” and “eNOS activation”, suggesting a relatively slow shutdown of the E/HE program in pre-HSCs. In contrast, subsets of E10.5 CD45^+^ IAC cells show lineage-specific differentiation bias; *Il7r* is up-regulated in 26%, and *Gata1* in 3% of E10.5 CD45^+^ IAC cells (Fig. 7f).

Previous scRNA-seq studies identified committed progenitors in E10.5 and E11.5 IACs but concluded they were contaminating erythro-myeloid progenitors (EMPs), likely originating from the yolk sac, that had been circulating in the blood and became attached to the IACs (Baron et al., 2018; Zhou et al., 2016). We addressed the possibility that the E10.5 CD45^+^ IAC cells we profiled are contaminating YS EMPs. Direct comparison shows E10.5 CD45^+^ IAC cells and YS EMP are molecularly and functionally distinct (Fig. 7h, Supplementary Fig. 7c). E10.5 CD45^+^ IAC cells contained progenitors of macrophages and granulocytes/monocytes, but very few erythroid or megakaryocytic progenitors compared to E9.5 YS EMPs (Fig. 7h). E10.5 CD45^+^ IACs have potent lymphoid potential; limiting dilution assays revealed a high frequency of cells (1:6) capable of producing B cells following culture on OP9 stromal cells or T cells on OP9 expressing the Notch ligand delta-like 1 (Fig. 7i,j). E10.5 CD45^-^ IAC cells also contained progenitors with lymphoid and myeloid potential, although their frequency was lower than in the E10.5 CD45^+^ IAC population. In summary, E10.5 CD45^+^ IAC cells represent a distinct wave of lympho-myeloid-biased progenitors in IACs that appear prior to E11.5 type II pre-HSCs.

## Discussion

In summary, our single cell analyses provide new insights into the process by which endothelial cells differentiate into pre-HSCs. First, we define a precursor of HE we have named pre-HE, in which multiple molecules and pathways known to regulate HSPC formation appear to act. Also, through trajectory analyses and genetic perturbation experiments we identified a bottleneck separating pre-HE from HE, indicative of a developmental barrier that must be overcome at that transition. It is long known that embryonic hematopoiesis is exquisitely sensitive to Runx1 dosage, as reduced Runx1 dosage decreases the number of HE cells, IACs and committed hematopoietic progenitors in the embryo (Cai et al., 2000; Lie et al., 2018; Wang et al., 1996). Our scRNA-Seq analyses show that the deficits caused by reduced Runx1 dosage are caused, at least in part, by the inefficient transition of pre-HE to HE cells. The molecular underpinnings of the bottleneck at the pre-HE to HE transition are not known. One possibility is that Runx1 expression may be actively repressed in the majority of pre-HE cells by TFs such as Sox17 (Bos et al., 2015; Lizama et al., 2015), which is highly expressed in pre-HE, and binding sites for which are accessible in pre-HE. A requirement for chromatin remodeling may also be a limiting factor in pre-HE, as multiple epigenetic regulatory proteins have been shown to affect Runx1 expression in HE, some of which may act at the pre-HE to HE transition (Kasper and Nicoli, 2018).

Prior to HE, Runx1 is expressed at low levels in a subset of endothelial cells, consistent with the chromatin accessibility of the P2 promoter and of several *Runx1* enhancers in endothelial cells. Runx1 expression in endothelial cells appears to be stochastic; it then becomes elevated in a subset of pre-HE cells, and is uniformly high in HE and IACs. The mechanism by which Runx1 expression is activated in a subset of pre-HE cells is not known, but our experiments provide some clues. scATAC-Seq revealed a distal enhancer in *Runx1*(−371), conserved in mammals, that first becomes accessible in pre-HE. Highly conserved TF motifs in the −371 enhancer include GATA, STAT, and JUN, implying that TFs that bind these motifs play a role in opening the enhancer in pre-HE. Gata2 expression is activated in a pulsatile manner in endothelial cells in the DA (Eich et al., 2018), which may contribute to the stochastic expression of Runx1 in arterial endothelial cells. STAT and JUN are recognized by TFs that are effectors of inflammatory signaling pathways, including type I and II interferons, and tumor necrosis factor, all of which promote HSPC formation from arterial endothelium (Espin-Palazon et al., 2014; Li et al., 2014; Sawamiphak et al., 2014). Hence well-known signaling pathways known to promote later *Runx1* expression in HE likely initiate Runx1 expression in a subset of pre-HE cells by activating the −371 enhancer. At later stages, in IACs and FL-HSCs, multiple additional enhancers, including the +23 enhancer gain accessibility and interact with the P1 promoter to further elevate Runx1 expression.

A second important concept gleaned from our data is that the IACs contain at least two distinct HSPC subtypes, committed lympho-myeloid biased progenitors and pre-HSCs, that can be distinguished molecularly. These appear sequentially, with CD45^+^ lympho-myeloid biased progenitors preceding the formation of type II pre-HSCs. The mechanisms underlying the generation of these two types of HSPCs is of great interest. It is not known, for example, if they independently differentiate from an equivalent population of immature IAC cells. Alternatively, they may be derived from distinct populations of HE cells. The lympho-myeloid biased progenitors are more developmentally “mature” compared to the type II pre-HSCs, suggesting that they are more driven towards terminal differentiation. A similar population of lympho-myeloid restricted progenitors that originates in the yolk sac colonizes the FL and thymus prior to HSCs (Boiers et al., 2013; Luis et al., 2016). Lympho-myeloid-biased progenitors in the arterial IACs may serve a similar function.

The earlier emergence of lympho-myeloid-biased progenitors in the arteries may have implications for ongoing efforts to generate pre-HSCs from ES cells. The acquisition of lymphoid potential is often used as a surrogate for pre-HSC formation. However, it is possible that conditions favoring the production of this earlier population of committed lympho-myeloid progenitors may be suboptimal for the later formation of pre-HSCs. If this is the case, then inhibiting the differentiation of lympho-myeloid progenitors in ES cell cultures may improve pre-HSC production ex vivo.

Data and analysis from this study can be interactively explored using a single cell EHT data explorer based on VisCello (Packer et al., 2019) and R shiny (Chang et al., 2019) (https://github.com/qinzhu/VisCello.eht).

## Data availability

Data generated during this study have been deposited in the Gene Expression Omnibus (GEO) with the accession number GSE137117. The EHT atlas can be interactively explored using VisCello (Packer et al., 2019) (https://github.com/qinzhu/VisCello.eht).

## Acknowledgments

This work was supported by National Institutes of Health grants R01HL091724 and R21AI133261 (NAS), R01HG006130 and R01 GM108716 (KT), R01HL136024 (KT and NAS), and under a Grant with the Pennsylvania Department of Health (KT and NAS). The Department specifically disclaims responsibility for any analyses, interpretations or conclusions. LB is supported by T32HL007439 and MM by T32DK007780. The Runx1-GFP mice were kindly provided by James Downing. We thank Sumedha Bagga for editorial assistance and Youtao Lu and Junhyong Kim for suggestions on analyses.

## Author contributions

Qin Zhu conducted the bioinformatics analyses and wrote the manuscript, Peng Gao and Changya Chen performed the scRNA-seq and scATAC-seq experiments, Joanna Tober, Laura Bennett, Yan Li, and Melanie Mumau purified and functionally validated the cell populations, Yasin Uzun and Wenbao Yu conducted additional bioinformatics analyses, Nancy A. Speck and Kai Tan directed the project and wrote the manuscript.

## Online Methods

### Animal husbandry

B6C3F1/J 3-week old female mice were purchased from Jackson Laboratories (Stock no: 100010). Females were injected with 5 IU pregnant mare serum gonadotropin (Sigma) and 48 hours later with 5 IU human chorionic gonadotropin (Sigma). Females were immediately paired overnight with C57BL6/J male mice. Runx1:GFP^8^ (*Runx1^tm4Dow^*) homozygous male mice were mated to super-ovulated B6C3F1/J 3-week old female mice to generate embryos for purification of E and HE cells. Female B6C3F1/J mice were mated with male B6129SF1/J mice for isolating fetal liver HSCs. The morning post mating is considered embryonic day (E) 0.5. E9.5-E11.5 embryos were accurately staged at the time of harvest by counting somites. Embryos that showed abnormal development were discarded. Mice were handled according to protocols approved by the University of Pennsylvania’s Institutional Animal Care and Use Committee and housed in a specific-pathogen-free facility.

### Embryo dissection and FACS

Yolk sacs were removed from embryos, and vitelline vessels were retained with the embryonic portion. The head, cardiac and pulmonary regions, liver, digestive tube, tail and limb buds were removed. The remaining portion containing the aorta-gonad-mesonephros (AGM) region, portions of somite, umbilical and vitelline vessels were collected. E9.5 and E10.5 yolk sacs were collected for isolation of EMPs. Tissues were dissected in phosphate buffered saline (PBS)/10% Fetal Bovine Serum (FBS) and Penicillin/Streptomycin (Sigma), followed by dissociation in 0.125 Collagenase (Sigma) for 1 hour. Tissues were washed and filtered through a 40-micron filter and resuspended in antibody solution (Supplementary Table 4). Cells were sorted on either BD Influx, MoFlow Astrios EQ (Beckman), BD Jazz, or BD Aria, all equipped with a 100-micron nozzle, and run at a pressure of 17 psi with flow rates less than 4000 events/second. Sorted cells for functional assays were collected in PBS/20% FBS/25mM HEPES. For scRNA-seq and scATAC-seq, cells were collected in IMDM/20% FBS in low-retention microcentrifuge tubes (Denville).

### OP9 co-culture assays

FACS sorted cells were plated in limiting dilutions on OP9 (ATCC) or OP9-delta-like 1 stromal cells in 96-well plates containing Minimum Essential Medium Eagle - alpha modification (alpha MEM), 20% FBS (Hyclone, Gibco) and Pen/Strep. 5 ng/mL Flt3L and 10 ng/mL IL-7 were added to the medium for OP9 co-cultures however, the medium for the OP9-DL1 co-cultures was supplemented with 5 ng/mL Flt3L and 1 ng/mL IL-7. Co-cultures were conducted for 10-13 days and subsequently, flow cytometry was performed on a LSR-II (BD). The flow cytometry antibody panel for OP9 co-cultures included the hematopoietic markers Mac1, Gr1, CD19, B220, and CD45, while the OP9-DL1 co-cultures were analyzed for CD45, CD90, and CD25. Limiting range was determined using the extreme limiting dilution analysis (ELDA) software analysis tool (Hu and Smyth, 2009).

### Methylcellulose assays

To enumerate erythroid, myeloid, and megakaryocyte progenitors, sorted cells were cultured in M3434 (StemCell Technologies) for 7 days. Colonies were scored based on morphological criteria.

### Hemogenic endothelial (HE) cell assay

Sorted cells were plated in limiting dilutions with OP9 stromal cells for 8-10 days in alpha MEM containing 10ng/mL of IL-3, IL-7, Flt3, and SCF. Cells were analyzed by flow cytometry for hematopoietic markers (B220, CD19, Mac1, Gr1, and CD45), and ELDA software analysis tool^39^ was used to determine the frequency of HE.

### scRNA-Seq

Sorted cells were immediately processed for library preparation using the 10x Genomics Chromium controller, in conjunction with the Chromium Single Cell 3′ Reagent Kit v2, according to manufacturer’s protocol (CG00052, Rev D). Libraries were quantified using the dsDNA High-Sensitivity (HS) Assay Kit (Invitrogen) on the Qubit fluorometer and the qPCR-based KAPA assay (Kapa Biosystems). Library quality assessment was performed on the Agilent 2100 Bioanalyzer in combination with the Agilent High Sensitivity DNA kit. Indexed libraries were pooled and sequenced on an Illumina HiSeq 4000 or NextSeq 550 using paired-end 26 × 98 bp read length.

### scATAC-Seq

Sorted cells were centrifuged at 300x *g* for 5 minutes (min) at 4 °C and resuspended in 50 µL of 1x PBS + 0.04% BSA. 45 µL of supernatant was carefully discarded, and 45 µL of chilled lysis buffer was added and mixed by pipetting gently. After incubation for 5 min on ice, 50 µL of chilled wash buffer was added without mixing. The mix was centrifuged at 500x *g* for 5 min at 4 °C, and 95 µL of supernatant was discarded. 45 µL of chilled diluted nuclei buffer was added without mixing, and the mix was centrifuged at 500x *g* for 5 min at 4 °C. The nuclei pellet was resuspended in 7 µL of chilled diluted nuclei buffer. 2 µL of nuclei suspension was used to determine the cell concentration by a Countess II cell counter (Invitrogen), and the remaining 5 µL of nuclei suspension was processed for library preparation using the Chromium Single Cell ATAC Reagent Kits protocol. Libraries were quantified using the dsDNA HS Assay Kit on the Qubit fluorometer and the qPCR-based KAPA assay. Library quality assessment was performed using the Agilent 2100 Bioanalyzer with the Agilent High Sensitivity DNA kit. Indexed libraries were pooled and sequenced on an Illumina NextSeq 550 using paired-end 50 × 50 bp read length.

### scRNA-Seq data analysis

#### Data pre-processing and filtering of non-endothelial and non-hematopoietic cells

Raw sequencing reads were first pre-processed with 10x Genomics Cell Ranger pipeline and aligned to the mouse mm10 reference genome. An initial filtering was performed on the raw gene-barcode matrix output by the Cell Ranger *cellranger count* function, removing barcodes that have less than 1000 transcripts (quantified by unique molecular identifier (UMI)) and 1000 expressed genes (“expressed” means that there is at least 1 transcript from the gene in the cell). Barcodes that pass this filter were considered as cells and were fed into downstream dimension reduction and clustering analysis. In the global UMAP with 37,766 cells combined from all datasets (Supplementary Fig. 2a), we noticed several contaminant cell types, including mesenchymal-like cells that express high levels of collagen, erythroid progenitors, and *Lyve1*^+^ endothelial cells that likely have a lymphatic or YS origin (Supplementary Fig. 2a). Since these contaminant cell types are not directly associated with EHT, we removed them from our downstream analyses, thereby obtaining a UMAP exclusively with endothelial cells and hematopoietic cells (Supplementary Fig. 2b).

Unsupervised clustering on the cleaned global UMAP reveals a clear separation between IACs and other hematopoietic progenitors. For example, Haptoglobin (*Hp*) is highly expressed in most YS EMPs, but has almost zero expression in IAC cells (Supplementary Fig. 2b). The proximity of EMP and IAC is likely driven by their shared differentiation potential, as both populations contain cells expressing *Gata1*, which mediates megakaryocyte differentiation (Ferreira et al., 2005) (Supplementary Fig. 2b). We found a *Bnip3^hi^* population in the E10.5 CD44^+^ E+HE+IAC samples, and a group of low-quality endothelial cells marked by low UMI counts per cell (Supplementary Fig. 2b). After filtering out these cells, we obtained a final UMAP with 23,081 cells representing the EHT trajectory (Supplementary Fig. 2c). We ran Louvain clustering on this global UMAP and assigned cell types based on differentially expressed genes (Supplementary Fig. 2c). The cell distributions of each dataset post-cleaning are shown in Fig. 2c.

#### Feature selection, dimension reduction, and unsupervised clustering

Gene-barcode UMI count matrix combined from all datasets was first processed with a standard pipeline utilizing the Monocle 3 package (Qiu et al., 2017). An initial variable expressed gene (VEG) selection was performed on the size-factor corrected, log2 transformed expression matrix using the feature dispersion table output by Monocle *estimate Dispersions* and *dispersionTable* function, which requires a gene to have *dispersion_empirical*/*dispersion_fit* > 0.5 and express at least more than 1 transcript in a minimum of 10 cells in order to be used as a VEG for dimension reduction. To produce a low dimensional embedding of the data, principal component analysis (PCA) was performed on the VEG-cell matrix, and the top PCs were used as features for the UMAP algorithm. UMAP was computed using the *umap* function in the uwot R package, with “cosine” distance metric, 20 nearest neighbors, and the rest of the parameters utilized were default. Louvain clustering was run on the k-nearest neighbor graph (k = 20) constructed from cell embeddings on the UMAP.

We noticed that VEGs selected using the Monocle 3 workflow contained genes that are cell-type specific, as well as genes associated with cell cycle and batch differences. Some highly expressed house-keeping genes are also called as VEGs, likely due to variation in sequencing depth across batches. The UMAP produced by the Monocle 3 pipeline is globally reflective of cell type, but locally affected by batch and cell cycle, causing some Louvain clusters to be less representative of underlying cell states.

To select features that are most reflective of cell type transitions during EHT and less affected by batch or cell cycle difference, we devised a feature selection procedure called informative feature (IFF) selection. IFF takes an initial clustering generated by other single-cell clustering methods such as Monocle 3 or SC3 (Kiselev et al., 2017) and then computes the expressed proportion of each gene for each cluster. Genes that are detected in less than 10% of cells in each cluster are filtered out. To determine the “inequality” of the gene’s expression across clusters, we calculated the expressed (non-zero) fraction for the remaining genes in each cluster and then determined Gini coefficient on the per-cluster-expressed-fraction vector. The distribution of Gini coefficients shows a clear peak on the left (Supplementary Fig. 5a). Genes in the peak are highly enriched in house-keeping and cell cycle function, while genes in the right long tail are strongly enriched with cell type specific ontologies (Supplementary Fig. 5b,c). This allows us to separate the majority of “cell-type-informative-features” from “ubiquitous features”. We found that IFF significantly improves the initial clustering result and identifies more underlying cell subtypes and states. The dimension reduction and clustering result based on IFF is highly robust across various PC inputs and parameter choices (UMAP with different parameters can be viewed by using the VisCello.eht package).

The IFF selection procedure is conceptually similar to the dpFeature selection introduced by the Monocle 2 package, which requires an initial clustering that is most reflective of cell type and selects cell-type-specific features by differential gene expression test. However, unlike the dpFeature, IFF selection is more permissive, as it does not require a gene to be significantly differentially expressed in one cell type to be selected. Additionally, genes expressed in a subset of clusters with a relatively weak but specific pattern are informative of cell type segregation, and a high Gini coefficient enables them to be selected as IFFs. Since Gini coefficient computation was based on the per-cluster-expressed-fraction rather than normalized expression, this procedure is also insensitive to various modeling assumptions underlying different single cell data normalization methods.

### Differential expression analysis

We ran differential expression analysis using the “sSeq” algorithm implemented in the cellrangerRkit package and used FDR < 0.05 and log2 fold change >1 to call differentially expressed genes (DEGs).

### Pathway enrichment analysis

To directly compute a per-cell enrichment score for each pathway in the Reactome database (Fabregat et al., 2016), we used an approach based on the AUCell package (Aibar et al., 2017). We slightly modified the standard AUCell pipeline; instead of using all genes for ranking, we initially removed the majority of housekeeping genes using the IFF selection method described above, thereby retaining genes that are mostly cell-type specific (top 25% of genes ranked by Gini coefficient). To derive pathways that are differentially active along the EHT trajectory and between pre-HSC and lympho-myeloid-biased progenitors, we subsequently performed stage-wise Student *t* test on the enrichment score (q-value <= 0.01, Supplementary Table 2 and 3). For Fig. 3f, 7g, Supplementary Fig. 3a, we removed pathways with fewer than 5 genes. Redundant pathways were removed if the Jaccard index (*number of shared genes/number of all genes)* for the pair of pathways was greater than 0.1 and the pathway had a higher q-value. For Supplementary Fig. 5b,c, we computed gene ontology (GO) enrichment using the ClusterProfiler package (Yu et al., 2012), q-value cutoff of 0.05 and ontology type “Biological Process” (BP).

### Pseudotime assignment

We applied Slingshot, one of the best performing trajectory inference methods based on a benchmark study of 45 methods (Saelens et al., 2019), to the cleaned data, as described above. Slingshot infers trajectory by fitting a principal curve along a user-selected low dimensional embedding of the data and assigns each cell a pseudotime based on its projection onto the curve. We used the UMAP in Fig. 2b, excluding FL-HSC, as the input to the Slingshot algorithm. The starting cluster was set to the “E (other)” population, as it includes the majority of E9.5 E, and the terminal cluster was set to the “IAC” population. Computed pseudotime was used for ordering cells along heatmaps in Fig. 2f, 3f, and Supplementary Fig. 3a. Cells in heatmaps of Fig. 7e,g and Supplementary Fig. 3b were ordered based on the PC score, rather than Slingshot-assigned pseudotime.

### Velocyto analysis

We used the “*velocyto run10x*” command with mm10 reference genome to quantify spliced and unspliced mRNAs. The output loom file was analyzed using the “velocyto.R” package. For the E10.5 E+HE+IAC dataset shown in Fig. 4, we sequenced ∼56 k reads/cell, and 13.3% UMIs contained unspliced intronic sequences. Velocyto analysis allows estimation of RNA velocity of single cells by distinguishing between unspliced and spliced mRNAs, which is predictive of the rate of transcriptome change along the EHT trajectory. We used the “*gene.relative.velocity.estimates*” function with *fit.quantile* = 0.05, *deltaT=1*, *kCells=20* to calculate RNA velocity and subsequently, visualized the velocity vector field in the UMAP using the “*show.velocity.on.embedding.cor*” function with 20-cell neighborhood and 80 grid points along each UMAP axis.

### Comparison with published scRNA-Seq data

We compared our data to three published scRNA-Seq datasets. Zhou *et al*. (Zhou et al., 2016) sequenced 181 cells including E, pre-HSC and HSC cells. Baron et al. (Baron et al., 2018) sequenced 1121 E, HE, EHT and IAC cells from E10 and E11 AGM using CEL-Seq. Mass *et al*. (Mass et al., 2016) sequenced ∼90 E10.25 EMP cells. First, we performed PCA on our data using shared genes with the public data, then used the top 10 PCs to compute a UMAP using the *umap* function from uwot package. Using the PCA loading matrix, we projected public data onto the same PCA space, then predicted UMAP embedding using *umap_transform* function with previously computed UMAP model. The final co-embedding for public data with our data are shown in Supplementary Fig. 7.

### scATAC-Seq data analysis

#### Data pre-processing and peak calling

scATAC-Seq reads were aligned to the mouse mm10 reference genome using the “*cellranger-atac mkfastq*” command. Peaks were called using MACS2 with the FDR cutoff of 0.10 and the following parameters: -q 0.10 --broad --broad-cutoff 0.10) (Zhang et al., 2008). Quality control statistics of the data are shown in Supplementary Fig. 4. We implemented a custom PERL script to quantify the reads overlapping with peaks individually for each cell. The read is considered to overlap with the peak if at least half of the read overlaps. By quantifying the reads for each individual cell, we obtained a peak-barcode matrix. The peak-barcode matrix underwent an initial filtering, requiring a barcode to have at least 2000 fragments, 1000 detected peaks and at least 20 percent fragments in peak to be considered as a “cell”. Additionally, for each peak, we computed the fraction of cells with non-zero value, and removed peaks that were detected in fewer than 1% of all cells. The final cleaned matrix contains 1670 cells and 150,427 peaks. We also computed a normalized data matrix by first log2 transforming the data (with 1 pseudocount added) and regressing out variance explained by total detected peaks per cell estimated with the “*lmFit*” function in the limma package (Ritchie et al., 2015). This normalized matrix was used for differential accessibility test and preliminary matching of scRNA-Seq and scATAC-Seq clusters.

#### scATAC-Seq clustering and differential accessibility analysis

Traditional feature selection methods designed for scRNA-Seq data do not work well on scATAC-Seq data, due to much greater sparsity and a close-to-binary data distribution (per genomic locus per cell, the expected read count is 0, 1 or 2). This makes it difficult to select the most informative features for clustering and cell identity mapping.

Using the IFF selection method described above, we were able to obtain a UMAP with cells separated into several distinct neighborhoods, which significantly improved the Louvain clustering quality (Supplementary Fig. 5d-f). To identify differentially accessible peaks (DAPs), we first binarized data as either open (>1) or closed (0), then calculated the fraction of cells that have open states for each peak in each cluster. We ran one-vs-rest Chi-square test on the fractions and called cluster-specific peaks (DAPs) using FDR < 0.05 and absolute log2 fold change >1. Fold change is defined as the ratio of open fractions between the two groups.

We observed that DAPs and DEGs show strong co-enrichment patterns in genomic loci for certain pairs of scATAC-Seq and scRNA-Seq clusters (Supplementary Fig. 5g). Many of the DAPs are located near promoters of the matched DEG, but some are much more distant. By computing a co-enrichment value, we established an initial mapping between DAPs and DEGs (Supplementary Fig. 5g).

#### Seurat alignment and co-embedding of scATAC-Seq and scRNA-Seq cells

We used the Seurat alignment algorithm (Stuart et al., 2019) to co-embed scATAC-Seq and scRNA-Seq cells onto a single UMAP shown in Fig. 5a. We first matched each cell-type-specific DEGs with corresponding DAPs 200 kb up and downstream of the TSS, using the method described above, obtaining 12768 links between 2379 DEGs and 10126 DAPs. We then summed up DAP fragments for each DEG, obtaining a gene-peak correspondence matrix as the “gene activity matrix” for Seurat alignment. Transfer anchors were computed using the “*FindTransferAnchors*” function in Seurat, with dimension reduction method set to “cca” (canonical correlation analysis). scATAC-Seq cells were then transferred to scRNA-Seq reference using the “*TransferData*” function, using 15 nearest neighbors and PCA for computing the weighted correction vectors. Finally, scATAC-Seq and scRNA-Seq Seurat objects were merged using the “*merge.Seurat*” function, and joint UMAP was computed with top 20 PCs and 15 nearest neighbors. Cell type labels were transferred from scRNA to scATAC using Seurat. Contaminant cell types, including mesenchymal, *Lyve1^+^* E and *Bnip3^hi^* E were removed from both scRNA-Seq and scATAC-Seq data, and only cells involved in EHT were used to generate the UMAP shown in Fig. 5a.

#### Inference of enhancer-promoter (E-P) links

The DAP-DEG matching procedure could link cell-type-specific peaks to nearby cell-type-specific genes. However, this association required both differential expression and differential chromatin accessibility to be significant, potentially missing E-P links with weaker signal. Inspired by the Seurat alignment algorithm and eQTL inference method, we used linear regression on matched scATAC-Seq and scRNA-Seq meta cells to find gene-distal peaks (defined as enhancers) that have a chromatin accessibility pattern significantly correlated with a gene’s expression. First, for each scATAC-Seq cell, we paired it to its nearest scRNA-Seq neighbor in the joint UMAP, establishing links between 1186 scATAC-Seq cells and 659 scRNA-Seq cells. Note that not all scRNA-Seq cells were paired with a scATAC-Seq cell and some scRNA-Seq cells were paired to multiple scATAC-Seq cells, but the paired scRNA-Seq cells were uniformly distributed along the EHT trajectory.

To overcome the sparsity in scRNA-Seq and scATAC-Seq data, we expanded the paired scATAC-Seq and scRNA-Seq neighbors to paired scATAC-Seq and scRNA-Seq neighborhoods by pooling counts from 10 nearest neighbors. We normalized the pooled expression and accessibility by regressing out per-meta-cell total counts from log2 transformed data, followed by z-score transformation. For each expressed gene, we ran linear regression with its pooled expression against pooled accessibility peaks 200kb upstream and downstream of its TSS in the paired scATAC-Seq meta-cell. Links with Bonferroni corrected p-values < 0.01 and regression coefficients > 0.1 were considered significant and these called peaks are likely enhancers that contribute to the corresponding gene’s expression. E-P links for *Runx1* shown in Figure 6 were called separately, including additional peaks within 500kb upstream and downstream of *Runx1* P1. For genes with multiple promoters (defined as regions 2000 bp upstream and 500 bp downstream of each TSS), we ran a second regression using the promoter accessibility as dependent variable and each called peak as independent variable. This allowed us to link each called enhancer to a specific promoter. We computed “cis-regulatory-activity matrix” based on the called E-Ps, and observed consistent pattern between a gene’s expression and its cis-regulatory-activity score (Supplementary Fig. 6).

#### TF activity assessment using chromVar

We assessed TF binding activity to enhancers with chromVar (Schep et al., 2017) using its default setting, but changed the default p.value cutoff in “*matchMotifs”* function to 0.1 / (2 * median(enhancer length)) to account for multiple testing. The input TF motifs were curated from the CIS-BP motif database (Weirauch et al., 2014). For each TF motif and each cell, a GC-bias and background-corrected deviation score was computed using the “*computeDeviations*” function, which represents the relative gain or loss of TF binding activity. Lastly, to identify TFs with stage-specific binding activity, we ran stage-wise Mann-Whitney U-test with the deviation scores, and considered those with FDR < 0.05 as significant.

